# Biophysical Attributes of CpG Presentation Control TLR9 Signaling to Differentially Polarize Systemic Immune-Responses

**DOI:** 10.1101/076935

**Authors:** Jardin A. Leleux, Pallab Pradhan, Krishnendu Roy

**Author notes:** Corresponding Author Krishnendu Roy, PhD, Robert A. Milton Chair, Director, Marcus Center for Cell-Therapy Characterization and Manufacturing (MC3M), Director, Center for ImmunoEngineering, The Wallace H. Coulter Department of Biomedical Engineering at Georgia Tech and Emory, The Parker H. Petit Institute for Bioengineering and Biosciences, Georgia Institute of Technology, Atlanta, GA.

## Abstract

It is currently unknown whether and how mammalian pathogen-recognition receptors (PRR) respond to biophysical patterns of pathogen-associated molecular danger-signals. Using synthetic pathogen-like particles (PLPs) that mimic physical properties of bacteria or large-viruses, we have discovered that the quality and quantity of Toll-like-receptor-9 (TLR9)-signaling by CpG in mouse dendritic cells (mDC) is uniquely dependent on biophysical attributes, specifically the surface-density of CpG and size of the presenting PLP. These physical patterns control DC-programming by regulating kinetics and magnitude of MyD88-IRAK4 signaling, NFκB-driven responses, and STAT3 phosphorylation, which in turn controls differential T cell responses and in vivo immune-polarization, especially T-helper 1 (Th1) versus T-helper 2 (Th2) antibody responses. Our findings suggest that innate immune cells can sense and respond not only to molecular, but also pathogen-associated physical patterns (PAPPs), broadening the tools for modulating immunity, helping to better understand innate response mechanisms to pathogens and develop new and improved vaccines.

## Introduction

Dendritic cells (DCs) are one of the most potent antigen presenting cells (APCs) in mammalian immune systems, with phenotypically, functionally and spatially unique subsets of DCs (Pulendran et al., 1999; Kadowaki et al., 2001; Colonna et al., 2004). DCs recognize and respond to pathogens of varying sizes, physical architectures, shapes and molecular compositions. Although many of the mechanisms of DC programming are still being elucidated, most research to date has focused on the biochemical composition of pathogen associated molecular patterns (PAMPs) and how they initiate various signaling pathways and immune polarization. It is well established that DCs elicit differential immune-phenotypes in response to various pathogens, which has been partially attributed to the type of PAMPs they encounter (Piccioli et al., 2009; Cervantes-Barragan et al., 2012). However, there is little knowledge on whether and how biophysical attributes of pathogens i.e. pathogen-associated physical patterns (PAPPs) affect DC signaling and subsequent immune responses.

PAMPs are fundamental in the design of next generation vaccines and immunotherapies as adjuvants to stimulate the immune system, though there are many limitations including unpredictable diffusion and undesired systemic toxicity (Hanson et al., 2015; Liu et al., 2014).

An alternative way of presenting vaccine components to immune cells is through particulate carriers, which are not only beneficial from a biomimetic point-of-view (pathogen-mimicking) but also could prevent retention unpredictability. Therefore, biomaterial-based, vaccine carriers designed to mimic pathogens (e.g. PLPs) are being widely investigated for delivery (Leleux and Roy, 2013; Ali et al., 2009; Stano et al., 2012; Ballester et al., 2015).

We hypothesized that variation of carrier physical properties will directly affect how PAMPs interact with pathogen recognition receptors (PRRs) in innate immune cells and modulate the resulting immune response (Lynn et al., 2015); an aspect that likely occurs in natural infections as well. Although several previous reports have indicated basic immune response differences when antigens and adjuvants are presented on carriers of various sizes (Fifis et al., 2004; Chen et al., 2011), the effect of TLR-ligand density and how the interplay of size and density influence PRR signaling in DCs and modulate immune responses has not been studied.

Here we investigated how fundamental biophysical patterns, mainly the density of PAMPs and the size of the PAMP-presenting entity, affect signaling in DCs, DC-subset tropism and downstream immunity. We used synthetic PLPs as a simple platform to mimic pathogen physical properties, allowing for better control of variables. Our results demonstrate that CpG-density and PLP-size controls DC signaling dynamics, resulting in unique cytokine signatures and T cell responses. Furthermore, we observed that skin DC subsets show unique preference towards PLPs of particular size, and that changes in physical patterns translate into unique immune responses with CpG-density and PLP-size differentially controlling antibody class switching. We went on to discover that PAPP-mediated immune-polarization is dictated by the unique relationship between MyD88, NFκB and STAT3 activation kinetics in DCs.

## Materials and Methods

### Animals

Six- to eight week old C57BL/6 mice (Jackson Labs) were used. All animal studies were approved by the Georgia Tech Institutional Animal Care and Use Committee.

### Materials

PLGA (Resomer 502H) was purchased from Sigma Aldrich. Polyethylenimine was from Polysciences Inc. (Warrington, PA). Culture materials were from Fisher. Antibodies were from Ebioscience (San Diego, CA), Biolegend (San Diego, CA) or LifeSpan Biosciences (Seattle, WA). ELISA kits were from Ebioscience. Unmodified CpG was from OligoFactory (Holliston, MA) and was evaluated for the endotoxin levels, to ensure endotoxin-free, highly pure CpG. Fluorescent CpG was from Integrated DNA Technologies (Coralville, IA). Raw-Blue reporter cells and all associated reagents were from Invivogen (San Diego).

### Synthesis of PEI modified PLGA micro- and nanoparticles

PLGA microparticles were prepared using a water-oil-water double emulsion, solvent evaporation method as previously published (Kasturi et al., 2005; Singh et al., 2008; Pradhan et al., 2014) PLGA nanoparticles were fabricated using a similar method with slight modifications.

### Particle characterization

Particle size and zeta potential was analyzed using dynamic light scattering (Malvern Zetasizer Nano). Loading of protein was quantified by analyzing supernatant fractions using a Micro-BCA assay (ThermoFisher). Loading of nucleic acids was quantified either using absorbance measurements or using a Ribogreen assay (Life Technologies). Fluorescent images of dual-loaded nanoparticles were taken using a Zeiss SIM microscope.

### Isolation of DC subsets

Murine DC subsets were generated using bone marrow derived in vitro culture systems (C57BL/6 mice). Bone marrow cells were cultured for 7 days in RPMI medium supplemented with 10% heat-inactivated Characterized FBS (HyClone), 1% penicillin/streptomycin, 1% 100mM sodium pyruvate, 1% non-essential amino acids and 0.5ml of 500mM 2-mercaptoethanol and supplemented with 20ng/ml GMCSF with 10ng/ml IL4 (Peprotech, Rocky Hill, NJ). Medium was replaced with supplemental growth factors on days 2, 4 and 6. On day 7, loosely adherent cells were collected.

### Flow cytometry for in vitro studies

Uptake of isolated DC subsets was analyzed using flow cytometry. PDL2+ were incubated with soluble or PEI-PLGA particle loaded APC- CpG (Integrated DNA Technologies, Coralville, IA) for 24 hours. CpG dosage was kept constant at 5ug/106 cells. To determine activation state of DCs, cells were incubated with PEI-PLGA particles loaded with CpG for 24 hours at a 5ug/106 cells. Cells were analyzed for fluorescent CpG uptake and supernatant was collected for secretome analysis.

### Antigen presentation of DCs to CD4+ T cells

Sorted CD4+ splenocytes (OTII) (using Miltenyi MACS kit) were cultured at a 2:1 ratio with activated DCs. Co-cultures were allowed to proceed for 72 hours after which supernatant was collected for ELISA assays.

### TLR9-IRAK4 Proximity Ligation Assay

BMDCs were sorted using a Pan DC Isolation kit (Miltenyi) to ensure purity. Cells were plated onto glass coverslips and PLP formulations were added for 30 mins or 6 hours. Cells were immediately fixed, blocked and stained with anti- TLR9 and anti-IRAK4 (LifeSpan Biosciences, Seattle, WA) overnight. The following day, cells were stained using the Duolink PLA system (Sigma, St. Louis, MO). Quantification of punctae was performed using Volocity software.

### NFkB and Activation Kinetics

The RAW-Blue reporter cell line (Invivogen, San Diego, CA) was used to determine CpG-dose dependent NFkB activation kinetics. Cells were cultured according to manufacturer’s instructions and treated with particles for 1, 6, 24, 48, 72 or 96 hours. Supernatant was then collected and added to the QUANTI-BLUE detection medium per the manufacturer’s instructions.

### Phospho-STAT3 Kinetics

Magnetic bead sorted (Pan DC Isolation Kit, Miltenyi) BMDCs were fixed, permeabilized and stained with anti-phospho-STAT3 (PY705) (BD Biosciences, NJ). Cells were then analyzed using an Accuri C6 flow cytometer (BD).

### Particle distribution and DC subset analysis of draining lymph nodes

Female C57BL/6 mice (5-6 weeks) were injected subcutaneously with 20ug of either soluble or particle-delivered Alexa-647-CpG. Control mice were given saline injections. Injections were performed either one day, five days or seven days prior to lymph node isolation. Mice were also given a booster injection on day 6 to evaluate the effect of boosting. Draining lymph nodes (inguinal) were isolated and processed into a single cell suspension or frozen for cryosectioning.

### Immunization protocol

Female C57BL/6 mice (5-6 weeks) were injected subcutaneously with either soluble or particle-delivered ovalbumin (10ug), and CpG (25ug). Ovalbumin and CpG were loaded onto the same particle for those groups. Control mice were given saline injections. Mice were given three injections at two week intervals. Lymphoid organs and blood were collected one week following the final injection. Cells were either stained or restimulated. Blood was centrifuged and serum collected for IgG titer analysis using ELISA.

### Lymphocyte restimulation

Lymphocytes isolated from the inguinal lymph nodes and spleen of immunized mice were restimulated with whole ovalbumin (200ug/ml) for 3 days. Supernatants were collected and analyzed using ELISA (Ebioscience).

### Lymph node and DC immunohistochemistry

Lymph nodes were frozen in OCT medium and cryosectioned at the Yerkes Pathology Center at Emory University. They were then blocked, stained and mounted. BMDCs were allowed to adhere to coverslips for 1 hour before treatment with particles. Coverslips were then washed, fixed, stained and mounted. All slides were imaged using either a Zeiss 700 Confocal microscope or a Perkin-Elmer Spinning Disk Confocal microscope.

## Results

### Quality and quantity of TLR9 activation is directly coupled to CpG density

Ligand density, defined as the average number of CpG molecules per unit area of the carrier-surface, is a fundamental parameter that could have profound effects on the quality and quantity of TLR9 signaling. We designed four formulations to simultaneously test the effects of size and CpG-density on activation of TLR9, especially the TLR9-MyD88-IRAK4 signaling pathway. We first fabricated cationic nano (~ 250 nm in average size) and microparticles (~1000 nm in average size) of poly(lactic-co-glycolic) acid, using our previously reported protocol (Kasturi et al., 2005; Pai Kasturi et al., 2006; Singh et al., 2008, 2009, 2011; Pradhan et al., 2014); thus allowing us to generate PLPs on which antigens and adjuvants of interest can be loaded together by electrostatic interactions (Table S1 and Figure S1a-b). Care was taken to ensure that the size distributions of micro and nanoparticles overlapped minimally (Fig. 1a). Note that the sizes of carriers were chosen to resemble that of large viruses (200-300 nm) or bacteria (1-2 microns). Particles were then electrostatically loaded with CpG, an anionic molecule. Both maximum loading (MPmax or NP_max_), and a lower loading level (MP_lo_ and NP_lo_) were used to formulate PLPs of high and low density and large and small sizes. Surface area of each particle was calculated to determine CpG density at maximum loading and loading density was optimized such that the low and maximum loaded formulations of each size of carriers had comparable CpG density (Fig. 1b, Table 1).

**Figure 1:**
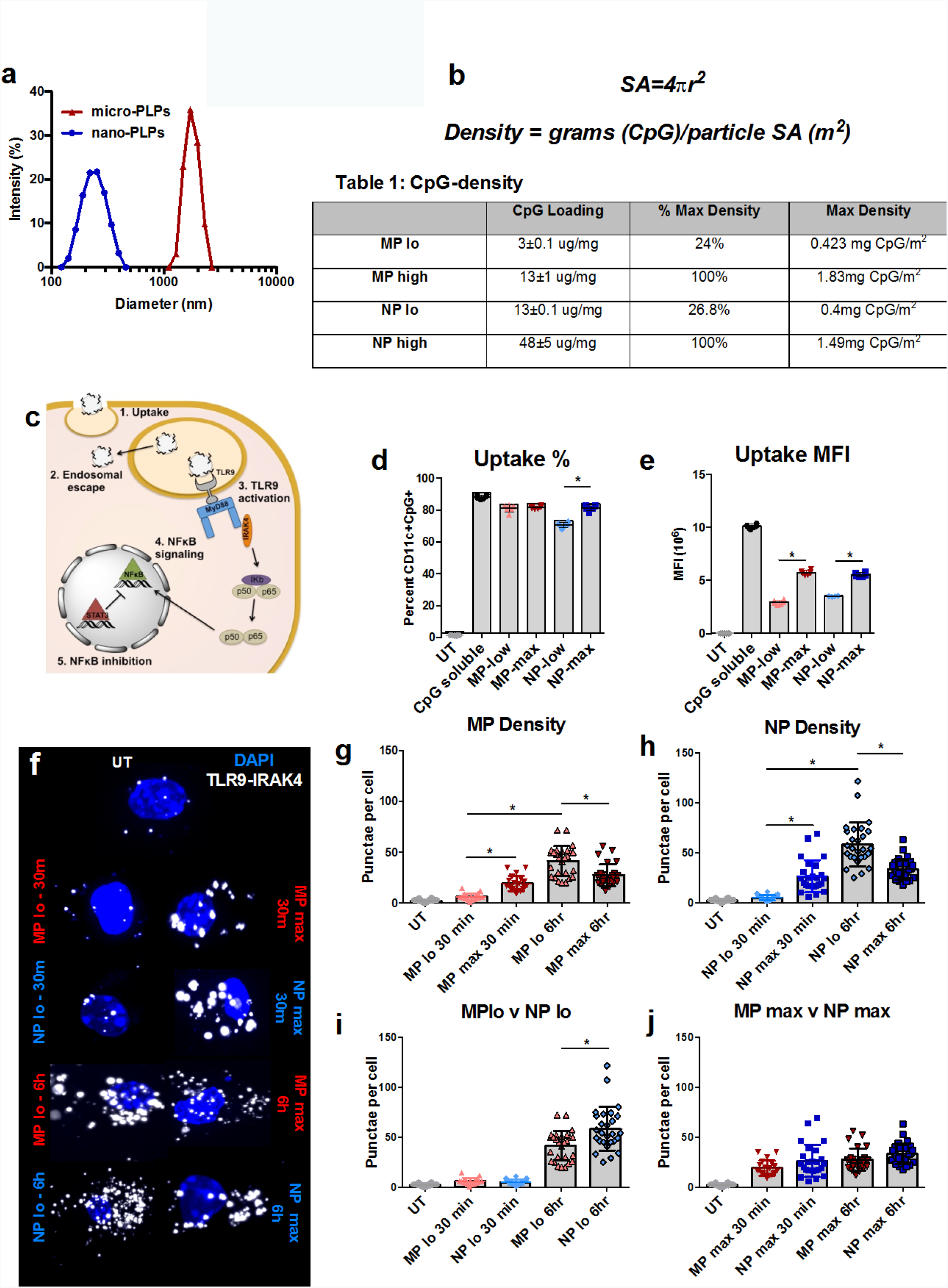
Particle size and CpG-ligand density have significant effects on uptake and TLR9 signaling kinetics in bone marrow derived dendritic cells. (A) Size distributions of micro- and nanoparticle formulations were measured using dynamic light scattering and have minimal overlap. (Table 1, B) Theoretical density was calculated using surface area formulas. CpG loading was confirmed by measuring the absorbance ratio (260/280nm) of loading supernatant. (C) CpG-TLR9 signaling has the potential to diverge at many points in the TLR9-NFkB axis. (D) Overall percentage of BMDCs that took up PLPs was similar across all formulations. (E) Max-density formulations promoted greater accumulation of CpG within individual cells than low-density formulations. (F) TLR9 activation quantified using a proximity ligation assay between TLR9 and IRAK4, where interactions are visualized as individual punctae. (G-H) CpG delivered on max-density PLPs induced rapid signaling of TLR9, within the first 30 minutes after delivery and was maintained for the first 6 hours of treatment. Signaling was significantly delayed for low-density PLPs, where TLR9 activity was not evident until 6 hours post-treatment. (I-J) Size did not seem to play a significant role in dictating signaling kinetics. Micro- or nano-PLP of corresponding densities did not signal differently with the exception of a small difference between signaling of low-density micro-PLPs and low-density nano-PLPs at 6 hours.

It is known that CpG ligation of endosomal TLR9 leads to activation and signal propagation, which can then promote transcription factor NFκB activity and induction of pro-inflammatory and/or anti-inflammatory programming of the exposed cell. To gain greater insight into PLP-driven DC programming, we investigated multiple points in the TLR9 - NFκB axis, including uptake, endosomal escape, TLR9 signaling through MyD88 to IRAK4, NFκB-mediated protein secretion and NFκB inhibition through STAT3 phosphorylation (an overview of the signaling cascade is shown in Fig. 1c). We observed that when mouse BMDCs were cultured with PLP formulations carrying fluorescent CpG for 24 hours, uptake varied with respect to CpG density. Mainly, MPMax and NP_Max_ formulations promoted accumulation of more CpG per cell (Fig. 1e), likely due to the increased mass ratio of CpG to PLGA in these formulations. The overall percentage of cells internalizing the CpG carriers remained similar (Fig. 1d).

Of PLPs that were taken up, the majority of the carriers remained co-localized with intracellular vesicle markers (i.e. EEA1 for early endosomes, LAMP1 for lysosomes and CD63 for general intracellular vesicles) indicating a similar degree of endosomal escape/retention across the various groups at early time points (Figs. S2a-c). To study TLR9/MyD88 signaling, we quantified the interactions between TLR9/MyD88 and IRAK4, a downstream signaling protein that is recruited after CpG engagement with an active TLR9 molecule (Figs. 1f-j). Signaling occurred within 30 minutes in cells treated with MP_max_ and continued 6 hours after initial treatment. However, signaling was significantly delayed in cells treated with low-density PLPs (NP_Max_), remaining similar to untreated cells at the early time point, but increasing strongly by 6 hours post-treatment (Fig. 1f). This was confirmed through image-quantification of puncta (Figs. 1g-h). Furthermore, we concluded that MyD88-mediated IRAK4 engagement was not a size-dependent phenomenon: signaling propagated with similar kinetics when density was matched (Figs. 1i-j). Finally, we also demonstrated that there is a density-dependent signaling magnitude difference at 6 hours, with the low-density groups promoting more robust signal at that time point (Figs.1f-g). There was also a slight increase in the magnitude of signaling for NP_lo_ formulations over MP_lo_ at this time point (Fig. 1h), but density played a larger role in this divergence.

### CpG-density and PLP-size control NF-kB driven protein production and STAT3 signaling

These results led us to investigate whether the observed delay in signaling resulted in differential downstream signaling and unique production and secretion of NFκB driven proteins. NFκB controls pro-inflammatory cytokine production as well as upregulation of costimulatory molecule expression, and therefore plays a critical role in DC maturation and polarization. Interestingly, despite a delay in TLR9 signaling, cells treated with MP_lo_ PLPs did experience rapid NFκB-driven protein secretion, equivalent to the MP_max_ formulations. At higher CpG doses (100ng-1ug), NFκB transcription events were evident in all groups, but were delayed in groups treated with NP_lo_ PLPs (Figs. 2c-d (insets), S3). Protein production converged after 48 hours of treatment, and cells treated with low density PLPs continued to ramp up their protein production even 4 days following initial treatment, while maximum density PLP-treated cells showed deceasing NFκB-driven protein production. These data indicated that for downstream NF-kB driven events, both size and density play a critical role.

It is known that CpG internalization also induces a regulatory response in parallel with pro-inflammatory signals to keep the immune response in check (Samarasinghe et al., 2006). STAT3 plays a major role in the inhibition of NFκB activity (Nefedova et al., 2005). We therefore measured phosphorylation of STAT3 over time in various PLP treated DCs. In line with other observations, STAT3 phosphorylation was quickly induced in low-density PLP formulations, particularly in NP_lo_ treated cells (Fig. 2e). It remained in a heightened state at 24 hours after treatment. This implies that the kinetics and quantity of STAT3 signaling is controlled by CpG density and PLP size; specifically STAT3 dominates signaling in NP_lo_ treated cells in early stages of DC programming, suppressing pro-inflammatory signaling and thus potentially promoting a phenotype consistent with regulatory DC development.

**Figure 2:**
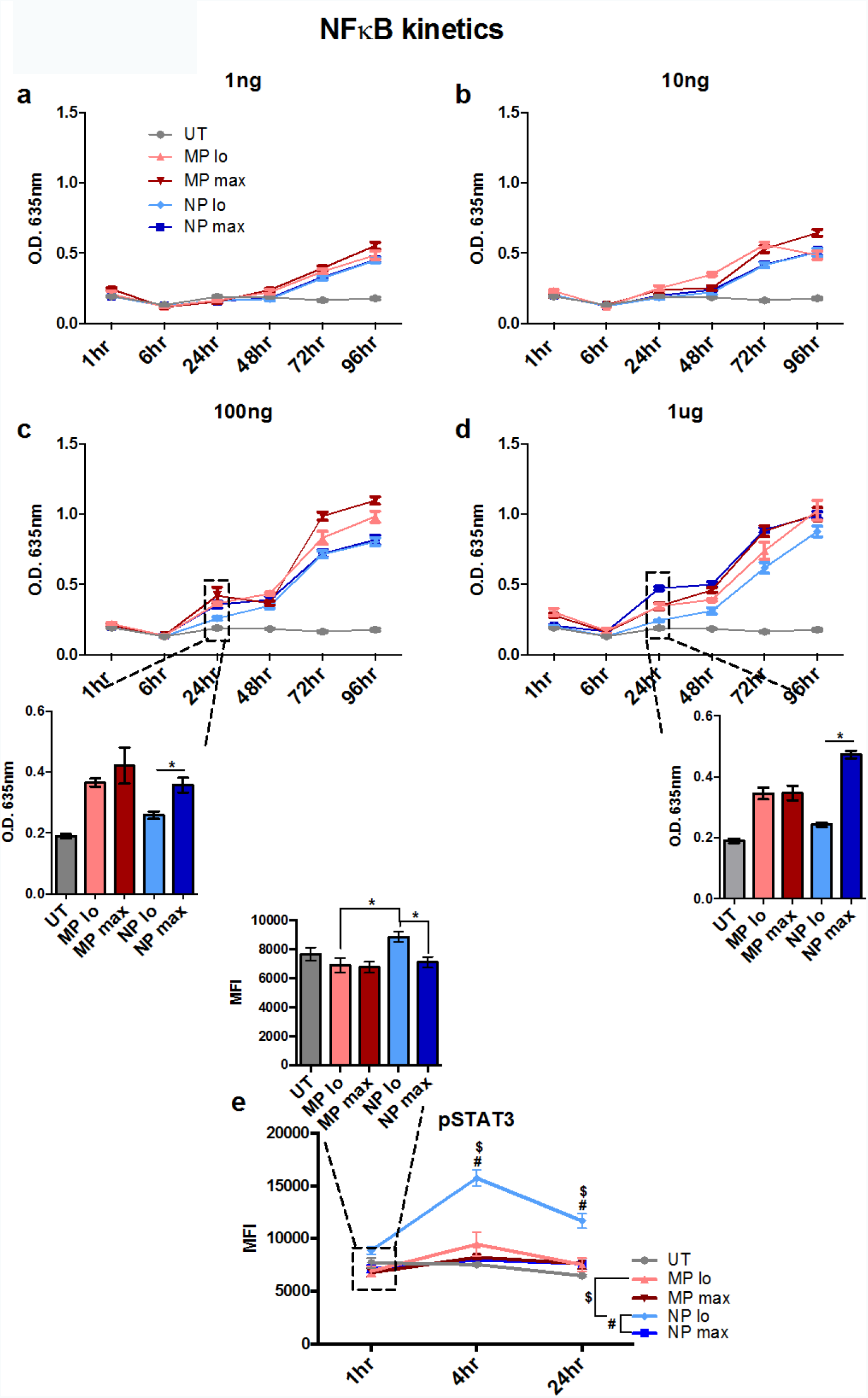
NFκB-mediated protein production is dampened and STAT3 activation induced by low-density nano-PLPs. (A-D) NFkB promotes the transcription of numerous DC pro-inflammatory signals. Despite the delay in TLR9 activation, low-density micro-PLPs induced NFkB dependent protein production, similar to max-density groups. While most other formulations also quickly induced transcription, low-density nano-PLPs induced a delayed response, which was not comparable to other treatment groups until 48 hours post-treatment. (E) Low-density nano-PLPs promote rapid phosphorylation of STAT3, an inhibitory factor responsible for IL10 production and NFkB regulation.

Formulation valency of surface-presented molecules has become another parameter of interested recently (Bennett et al., 2015). We calculated the valency of our formulations and determined that the valency of low density microparticle and high density nanoparticle were within the same order of magnitude (Supplementary Table 2.1). To determine whether valency played a major role, much like density, we statistically compared those groups in all of the data sets presented in this study (Supplementary Table 2.2). Less than half of our observations of these groups were significantly different, indicating that valency may play a role, but it is likely less influential than size or density.

### PLP-dependent signaling promotes diverging DC and T cell cytokine profiles

Immunomodulatory cytokine production (i.e. the DC secretome) is indicative of DC programming and provides predictive power for how DCs regulate T cell activation and subsequent systemic immune responses. We next investigated how CpG-density could be used to effectively tune DC secretome and subsequent T cell activity. Consistent with the rapid kinetics of TLR9 activation and NFκB activity, DCs treated with maximum density formulations were highly activated 24 hours following treatment, characterized by high levels of IL12p70, a highly pro-inflammatory cytokine (Fig. 3a). IL10 secretion was similar regardless of PLP treatment (Fig. 3b).

**Figure 3:**
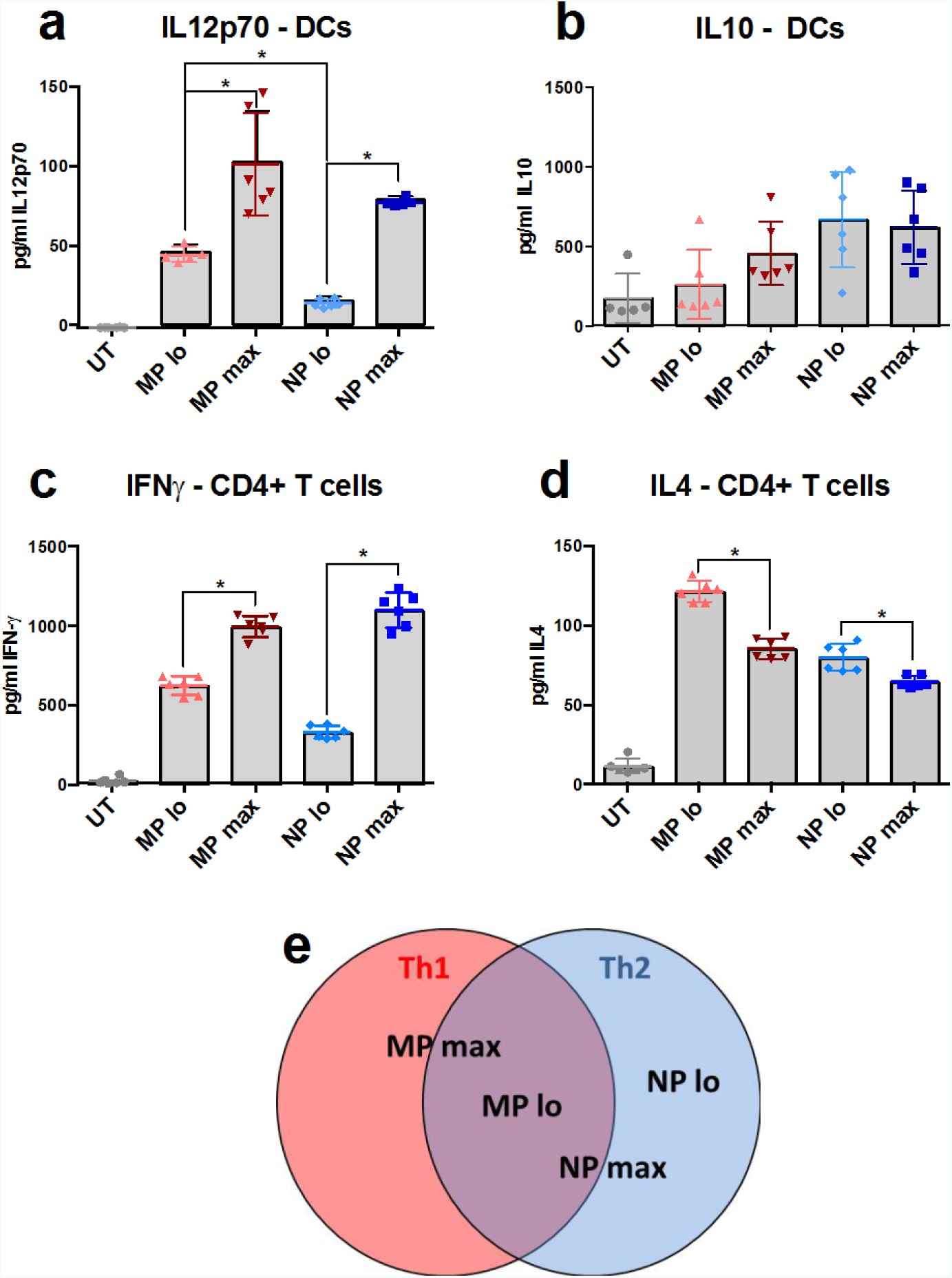
PLP-size and CpG-density influence DC and T cell cytokine profiles. (A) Dendritic cells were derived from bone marrow. Max-density PLPs promoted greater secretion of IL12p70 in activated BMDCs 24 hours after initial treatment. (B) IL10 production was low regardless of treatment 24 hours following initial treatment. (C) IFN-γ secretion patterns of CD4+ T cells that were co-cultured with activated BMDCs mirrored IL12p70 profiles; with max-density PLP treated DCs inducing greater production. (D) IL4 secretion patterns followed an inverse pattern, with low- density formulations inducing greater secretion of IL4 than their max-density counterparts. (E) Schematic describing the shift in systemic immune polarization observed in after treatment of each formulation. Data are representative of multiple experiments each with n=6. Error bars represent standard deviation.

Cytokine secretion of CD4+ antigen-specific T cells following co-culture with treated DCs in vitro revealed profiles that correlate with the potent activation associated with maximum density formulations. Mainly, we observed high levels of interferon-γ (IFN-γ) being produced in CD4+ antigen-specific T cells when cultured in the presence of maximum density PLP-treated DCs (Fig. 3c). IL4 secretion followed the inverse trend, with low density formulations promoting greater secretion than their maximum density counterparts (Fig. 3d). Experiments conducted with CD8+ cells revealed similar but dampened IFN-γ production trends and also did not show significant differences in IL4 production regardless of treatment (S4a-b). This begins to provide a framework for the important role that physical patterns (PAPP) play in controlling T-cell mediated systemic immunity (Fig. 3e).

### PLP size influences DC subset migration to draining lymph nodes

A significant body of recent work has investigated how size affects the ability of particulate formulations to traffic to the draining lymph nodes, either through passive lymphatic drainage or active phagocyte transport (Reddy et al., 2006; Swartz et al., 2008; Gerner et al., 2015); but little is known on which immune cell subtypes are targeted by particles of a specific size-range and how that differential tropism, if any, effects immune polarization. To confirm that both micro- and nano-PLPs traffic from skin to dLNs, we injected mice with PLP formulations loaded with fluorescent CpG oligonucleotide subcutaneously and collected draining (inguinal) lymph nodes 24 hours later. We observed that both micro and nano-PLPs were located in T cell zones (Fig. 4a left, right, respectively). Large amounts of micro-PLPs were also present in the periphery of the dLN and are likely associated with cells of the lymphatic sinuses, as described previously (Gerner et al., 2015). Overall, micro-PLPs facilitated greater trafficking of CpG to draining lymph nodes at 24 hours, either by migratory DCs or passive drainage (Fig 1b, inset). To further probe which migratory cell types are responsible for transporting PLPs from the injection site over time, we first analyzed DC (CD11c+) and macrophage (CD169+) compartments to confirm that DCs were responsible for a large portion of the PLP transport. We did find substantial quantities of CpG+ PLPs in both, as expected. Interestingly, there does seem to be a maximum density preference in both cell types (Fig 4b-c). We then stained cells isolated from dLNs for multiple DC subsets known to be present in the skin (Fig 4d). For these studies, lymph nodes were collected 1 day following the initial injection.

**Figure 4:**
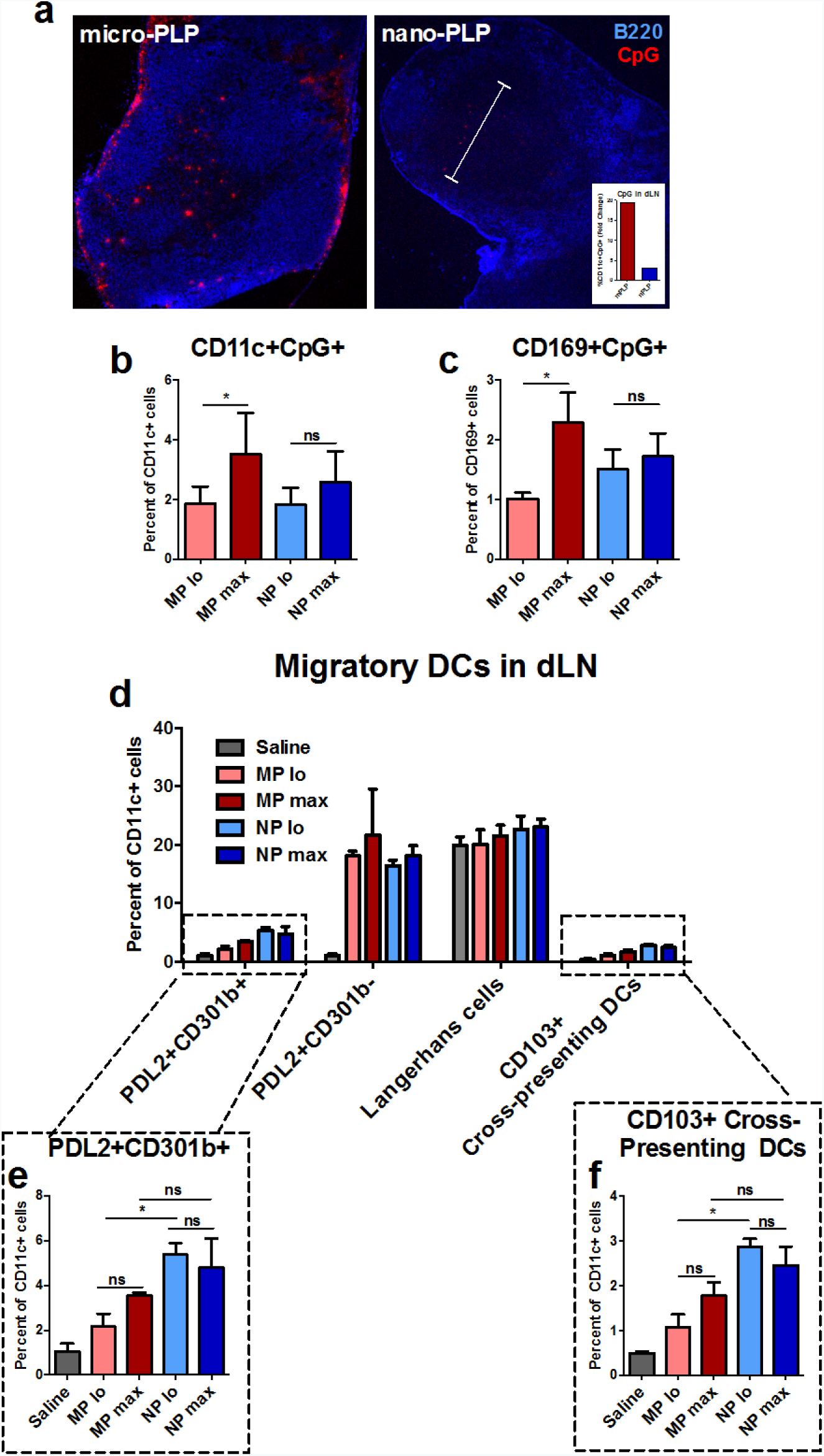
Migratory DCs exhibit PAPP-dependent tropism in vivo. (A) C57Bl/6 mice were injected with micro-PLP or nano-PLP formulations carrying fluorescent CpG. Draining lymph nodes were collected 24 hours later and cryosectioned to determine particle distribution. Slices were stained with anti-B220 to outline lymph node morphology. Inset graph shows average fold change of the percentage of CD11c+CpG+ cells present in dLNs 1 day after injection. (B-C) Draining lymph nodes of C57Bl/6 mice were collected 1 day post injection of fluorescent CpG formulations to evaluate migration of DCs or macrophages from injection site. Both transported PLP and exhibited density-dependent preferences. (D-F) DC subsets were evaluated in the dLN one day after initial injection with fluorescent CpG formulations. PDL2+CD301b+ dermal DCs and CD301+ cross-presenting dermal DCs both preferentially took up nano-PLP formulations. Data are representative of multiple independent experiments each with n=3. Error bars represent standard deviation.

These studies confirmed that some skin-resident DC subsets exhibit unique size-dependent preferences for PLPs. Mainly, PDL2+CD301b+ dermal DCs and CD103+ cross-presenting dermal DCs both exhibited a preference for nano-PLPs, particularly when comparing low-density formulations (Figs. 4e-f). Finally, we analyzed all DC subsets that were CpG+ to determine which subsets were responsible for transporting PLPs. PDL2+ dermal DCs made up over 85% of the CpG+ cells for all formulations (Fig. S4a). Interestingly, the BMDC culture system we have adopted promotes high expression of PDL2, further confirming the validity of our in vitro findings (Figs.4b-c).

### In vivo systemic immunity can be controlled by PAPPs

To identify whether physical patterns of CpG-carriers (CpG-density and PLP-size) can be utilized to modulate adaptive immunity polarization in vivo, we employed a vaccination model using ovalbumin (OVA) as the model antigen, which has been widely utilized to study immune responses (Moon et al., 2011; Hanson et al., 2015). In our in vitro studies, we showed that PAPPs play an essential role in directing DC-mediated CD4+ T cell behavior. Helper T cells provide signals to lymphoid resident B cells to promote class switching of secreted antibodies, which occurs in the germinal centers of lymphoid organs. Therefore, we first studied the functionality of activated B cells in vaccinated mice. Delivery of vaccine components on polymer PLPs increased the production of IgG1 over soluble vaccines (Fig. 5a, S5a), corroborating previous observations (Hanson et al., 2015). Strikingly, only mice that were treated with max-density formulations had high levels of IgG2c compared to low-density counterparts. IgG2c is a T-helper-1 (Th1) antibody isotype and is associated with a type 1 cytotoxic response (Fig. 5b-c) and anti-bacterial immunity. Furthermore, size seems to play a corresponding role in Th1 class switching, evidenced by statistically increased levels of IgG2c in mice treated with micro-formulations, independent of density (Fig. 5b-c, S5b-c). We also demonstrated increased germinal center formation and IgG production in inguinal lymph nodes of all treated groups by immunohistochemistry (Fig. 5d, S5d-e), indicating that all were immunologically active.

**Figure 5:**
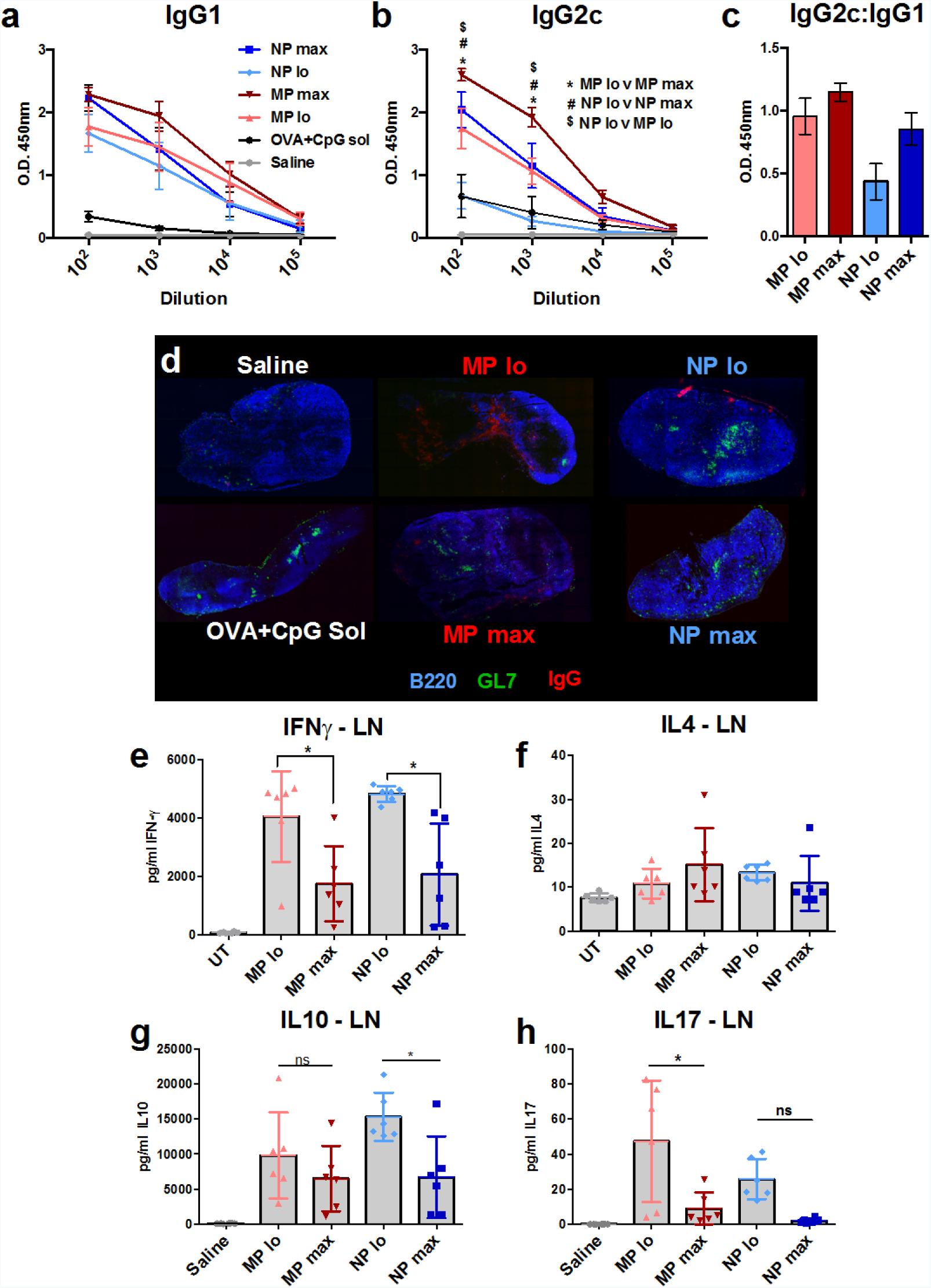
CpG-density and size skew systemic immune response. (A) Mice were inoculated with either soluble, micro- or nano-PLPs with either low- or max- density of CpG (20ug) simultaneously delivering co-loaded ovalbumin (10ug) 3 times at 2 week intervals. Lymphoid organs and serum were harvested one week after the final booster. IgG1 levels were higher in serum of all PLP groups, indicating a robust humoral response. (B-C) Max- density PLP vaccines, especially max- density micro-PLPs, induced class switching in activated B cells to produce significant levels of IgG2c, which indicates Th1 polarization of the cellular response. The ratio of IgG2c to IgG1 indicates a distinct shift in immune response polarization, particularly in nano-PLP treated mice. This is indicative of immunomodulatory control solely based on physical parameters of each formulation. (D) Immunohistochemistry confirms germinal center formation and class switching to IgG in all treatment groups. (E) Lymph node cells were harvested and restimulated with whole ovalbumin protein for 3 days. IFNγ production was high in groups treated with low- density PLPs at this time point. (F) IL4 remained low in all groups. (G) The anti-inflammatory cytokine IL10 was increased in mice treated with PLPs, especially in those given low- density nano-PLPs, indicating elevated regulatory activity in the LNs of these mice. (H) IL17 was marginally increased in low- density PLP groups, but overall was produced in low amounts. Data are representative of experiments each with n=6. Error bars represent standard error of the mean.

To confirm that the observed systemic response was indeed T cell mediated, we restimulated LN cells from vaccinated mice with whole OVA antigen and evaluated their secretome. We first characterized the CD4:CD8 ratio in dLNs and determined there is only a slight increase in the ratio in mice treated with low-density MPs (Fig. S6a). This may be correlated with a slight increase in regulatory T cells in the same mice (Fig. S6b). IFN-γ was produced in greater amounts in low density -PLP treated mice at this time point in both LN and splenic compartments (Figs. 5e, S7a, S8a). IL4 was not produced in very large amounts regardless of treatment (Figs 5f, S7b). Interestingly, LN (and spleen) lymphocytes of mice treated with NP_lo_- PLPs secreted the anti-inflammatory cytokine IL10 in greater quantities than NP_max_ or MP formulations, indicating higher regulatory activity (Figs. 5g, S8c, S8c). IL17 was preferentially secreted in mice that were treated with LV-PLPs (Figs. 5h, S7d). These observations clearly establish unique PAPP-driven immunological effects, confirming the importance of physical patterns in vaccine design.

## Discussion

T cell maturation is a hallmark of an effective immune response and is critical for both killing of infected and cancerous cells as well as building a potent humoral response (Jung et al., 2002). It has been appreciated for decades that DCs play an essential role in initiation of adaptive immune responses by presenting antigens and stimulatory signals to dictate subsequent T cell responses (Kapsenberg, 2003). Recent literature has expanded on the breadth of DC mediated responses, including functional characterization of many subsets (Guilliams et al., 2010), highlighting the importance of investigating how vaccine components interact with each of these subsets differently and what consequences this has for downstream immunity.

Biomaterials are currently being increasingly used in vaccine design to gain greater control of immunomodulatory effects, but the degree to which material properties, particularly adjuvant density, contribute to the efficacy and potency of the vaccine is not well understood(Elamanchili et al., 2004; Pradhan et al., 2014). While Bandyopadhyay et al. studied the effect of the targeting ligand DEC-205 density on DC uptake (Bandyopadhyay et al., 2011), there are no studies to our knowledge that expand this investigation to PAMP-density. Carrier size has been studied extensively, but there is neither a consensus on how size affects DC uptake and maturation uniquely by subset or whether particle size can be used to modulate peripheral DC-mediated T cell immunity. Moreover, there is a distinct lack of information regarding how DC signaling is affected by these parameters. We have shown that not only are PLPs taken up by different peripheral DC subsets in a size dependent manner but their migration to draining lymph nodes and T cell engagement also varies with size. All of these advances will educate the design of vaccines and immunotherapies to improve not only overall efficacy, but also direct immunity to the most appropriate adaptive response for a given indication.

Interestingly, our findings also suggest that PLPs that mimic bacteria or viruses in size (micro-PLPs and nano-PLPs, respectively) also promote responses that are similar to the corresponding pathogen. For example, T cells are driven to secrete IFNγ after bacterial infection, which has been shown to promote antibody class switching to IgG2c (Barr et al., 2009; Havell, 1982; Klinman et al., 1996). This allows bacterial cells to be targeted, tagged (with IgG2c antibodies) and cleared via antibody-dependent-cell-mediated cytotoxicity and natural killer cell activation (Catalona et al., 1981). On the other hand, viral infections promote the activation of CD103+ cross-presenting DCs and less antibody production (Lund et al., 2003; Mothe et al., 2002). These hallmarks align with the corresponding PLP-driven responses we’ve described in this study, validating the significance of the effects of this material property on the immune response.

We believe that micro-PLP’s (particularly the max-density formulation) ability to promote the production of an NFκB-mediated, pro-inflammatory DC cytokine profile could explain the Th1 skewing we observe systemically. This response is in line with expected results of a CpG oligonucleotide supplemented vaccine (Pradhan et al., 2014). The function of IL17 is still controversial but may be another mechanism by which micro-PLPs promote a response that mirrors one induced following bacterial infection (Jin and Dong, 2013).

Nano-PLPs, on the other hand, simultaneously skew the immune response towards an anti-viral like response, as well as a more defined regulatory response in DCs, driven by STAT3. We confirmed that low-density nano-PLPs promote STAT3-mediated induction of anti-inflammatory cytokine production and inhibition of NFκB activation. Multiple modes of activation have been described for STAT3, one being through the IL10-receptor binding of extracellular IL10 (Saraiva and O’Garra, 2010). However, based on the time scale of our observations, another mechanism must be dominant. We hypothesize that, in our system, STAT3 could be activated downstream of the PI3K pathway, which is independent of MyD88-IRAK4 activation and can be initiated very quickly after TLR activation (Brown et al., 2011; Gunzl et al., 2010).

Furthermore, it has been shown that activation of PI3K and downstream STAT3 both result a decrease in pro-inflammatory cytokine (i.e. IL12p70) production, similar to our observation (Vogt and Hart, 2011). Further studies need to be performed to determine whether this is one cause of the increased regulatory response observed in mice treated with low-density nano-PLPs. These novel findings enhance our understanding of the interaction between DCs and particulate vaccine carriers, as well as contributing to the control over immune modulation.

The finding that CpG-density plays a critical role in the size-driven dichotomy is established for the first time through this study. According to our observations, DC programming can be tuned by simply varying CpG-density, demonstrated here with the TLR9 agonist CpG. A recent study by Ohto et al. characterized the interaction of stimulatory CpG with TLR9, indicating that activated TLR9 dimerizes prior to signaling (Ohto et al., 2015). Other TLRs must also cluster for robust signal propagation (Inoue and Shinohara, 2014). Therefore, it is our hypothesis that, at low densities, the accessibility of CpG on PLPs may not be high enough to allow effective dimerization and or clustering of engaged TLR9 molecules. At later time points, only after significant accumulation of nano-PLPs (low density) in DC endosomes does CpG reach the threshold for proper signaling. This allows for other signaling mechanisms to take over and dictate DC programming.

Overall, our results provide new evidence that biophysical properties of CpG presentation, specifically ligand-density and size of presenting entity, have a distinct and profound effect on the kinetics and magnitude of MyD88, NF-kB and STAT3 signaling in DC, thereby directly modulating DC programming and the resulting adaptive immune response. This has significant implications in understanding pathogen-initiated responses in mammalian immune system and also for designing improved, precisely-tunable vaccines and immunomodulatory therapies against cancer, infectious diseases and autoimmune applications.

### Funding

This work was partially supported by the Georgia Tech Foundation, the Carol Ann and David D. Flanagan Fellowship, and the Robert A. Milton Fellowship to KR. JAL is a recipient of the National Science Foundation Graduate Research Fellowship.

## Acknowledgements

The authors thank Kristin Loomis and Prof. Philip J. Santangelo of Georgia Tech for protocol and guidance with the proximity ligation assays. The authors also thank Prof. Dmitry Shayakhmetov of Emory University for reading the manuscript and providing feedback. Additionally, the authors thank Virginia Bliss and Deepa Kodandera at the Yerkes Pathology Lab for their help with cryosectioning.

## Author contributions

JAL designed and executed experiments, analyzed data and wrote the manuscript. PP contributed to experimental execution and manuscript feedback. KR supervised experimental design and contributed significantly to manuscript preparation. All authors reviewed and approved final manuscript.

## References

Ali, O.A., N. Huebsch, L. Cao, G. Dranoff, and D.J. Mooney. 2009. Infection-mimicking materials to program dendritic cells in situ. Nat. Mater. 8:151–158. doi:10.1038/nmat2357.

Ballester, M., L. Jeanbart, A. de Titta, C. Nembrini, B.J. Marsland, J.A. Hubbell, and M.A. Swartz. 2015. Nanoparticle conjugation enhances the immunomodulatory effects of intranasally delivered CpG in house dust mite-allergic mice. Sci. Rep. 5:14274. doi:10.1038/srep14274.

Bandyopadhyay, A., R.L. Fine, S. Demento, L.K. Bockenstedt, and T.M. Fahmy. 2011. The impact of nanoparticle ligand density on dendritic-cell targeted vaccines. Biomaterials. 32:3094–3105. doi:10.1016/j.biomaterials.2010.12.054.

Barr, T.A., S. Brown, P. Mastroeni, and D. Gray. 2009. B Cell Intrinsic MyD88 Signals Drive IFN-Production from T Cells and Control Switching to IgG2c. J. Immunol. 183:1005–1012. doi:10.4049/jimmunol.0803706.

Bennett, N.R., D.B. Zwick, A.H. Courtney, and L.L. Kiessling. 2015. Multivalent Antigens for Promoting B and T Cell Activation. ACS Chem. Biol. 10:1817–1824. doi:10.1021/acschembio.5b00239.

Brown, J., H. Wang, G.N. Hajishengallis, and M. Martin. 2011. TLR-signaling Networks: An Integration of Adaptor Molecules, Kinases, and Cross-talk. J. Dent. Res. 90:417–427. doi:10.1177/0022034510381264.

Catalona, W.J., T.L. Ratliff, and R.E. McCool. 1981. γ Interferon induced by S. aureus protein A augments natural killing and ADCC. Nature. 291:77–79. doi:10.1038/291077a0.

Cervantes-Barragan, L., K.L. Lewis, S. Firner, V. Thiel, S. Hugues, W. Reith, B. Ludewig, and B. Reizis. 2012. Plasmacytoid dendritic cells control T-cell response to chronic viral infection. Proc. Natl. Acad. Sci. 109:3012–3017. doi:10.1073/pnas.1117359109.

Chen, H.C., B. Sun, K.K. Tran, and H. Shen. 2011. Effects of particle size on toll-like receptor 9-mediated cytokine profiles. Biomaterials. 32:1731–1737. doi:10.1016/j.biomaterials.2010.10.059.

Colonna, M., G. Trinchieri, and Y.-J. Liu. 2004. Plasmacytoid dendritic cells in immunity. Nat. Immunol. 5:1219–1226. doi:10.1038/ni1141.

Elamanchili, P., M. Diwan, M. Cao, and J. Samuel. 2004. Characterization of poly(d,l-lactic-co-glycolic acid) based nanoparticulate system for enhanced delivery of antigens to dendritic cells. Vaccine. 22:2406–2412. doi:10.1016/j.vaccine.2003.12.032.

Fifis, T., A. Gamvrellis, B. Crimeen-Irwin, G.A. Pietersz, J. Li, P.L. Mottram, I.F.C. McKenzie, and M. Plebanski. 2004. Size-Dependent Immunogenicity: Therapeutic and Protective Properties of Nano-Vaccines against Tumors. J. Immunol. 173:3148–3154. doi:10.4049/jimmunol.173.5.3148.

Gerner, M.Y., P. Torabi-Parizi, and R.N. Germain. 2015. Strategically localized dendritic cells promote rapid T cell responses to lymph-borne particulate antigens. Immunity. 42:172–185. doi:10.1016/j.immuni.2014.12.024.

Guilliams, M., S. Henri, S. Tamoutounour, L. Ardouin, I. Schwartz-Cornil, M. Dalod, and B. Malissen. 2010. From skin dendritic cells to a simplified classification of human and mouse dendritic cell subsets. Eur. J. Immunol. 40:2089–2094. doi:10.1002/eji.201040498.

Gunzl, P., K. Bauer, E. Hainzl, U. Matt, B. Dillinger, B. Mahr, S. Knapp, B.R. Binder, and G. Schabbauer. 2010. Anti-inflammatory properties of the PI3K pathway are mediated by IL-10/DUSP regulation. J. Leukoc. Biol. 88:1259–1269. doi:10.1189/jlb.0110001.

Hanson, M.C., M.P. Crespo, W. Abraham, K.D. Moynihan, G.L. Szeto, S.H. Chen, M.B. Melo, S. Mueller, and D.J. Irvine. 2015. Nanoparticulate STING agonists are potent lymph node–targeted vaccine adjuvants. J. Clin. Invest. 125:2532–2546. doi:10.1172/JCI79915.

Havell, E.A. 1982. Enhanced production of murine interferon gamma by T cells generated in response to bacterial infection. J. Exp. Med. 156:112–127. doi:10.1084/jem.156.1.112.

Inoue, M., and M.L. Shinohara. 2014. Clustering of Pattern Recognition Receptors for Fungal Detection. PLoS Pathog. 10:e1003873. doi:10.1371/journal.ppat.1003873.

Jin, W., and C. Dong. 2013. IL-17 cytokines in immunity and inflammation. Emerg. Microbes Infect. 2:e60. doi:10.1038/emi.2013.58.

Jung, S., D. Unutmaz, P. Wong, G.-I. Sano, K. De los Santos, T. Sparwasser, S. Wu, S. Vuthoori, K. Ko, F. Zavala, E.G. Pamer, D.R. Littman, and R.A. Lang. 2002. In Vivo Depletion of CD11c+ Dendritic Cells Abrogates Priming of CD8+ T Cells by Exogenous Cell-Associated Antigens. Immunity. 17:211–220. doi:10.1016/S1074-7613(02)00365-5.

Kadowaki, N., S. Ho, S. Antonenko, R. de Waal Malefyt, R.A. Kastelein, F. Bazan, and Y.-J. Liu. 2001. Subsets of Human Dendritic Cell Precursors Express Different Toll-like Receptors and Respond to Different Microbial Antigens. J. Exp. Med. 194:863–870. doi:10.1084/jem.194.6.863.

Kapsenberg, M.L. 2003. Dendritic-cell control of pathogen-driven T-cell polarization. Nat. Rev. Immunol. 3:984–993. doi:10.1038/nri1246.

Kasturi, S.P., K. Sachaphibulkij, and K. Roy. 2005. Covalent conjugation of polyethyleneimine on biodegradable microparticles for delivery of plasmid DNA vaccines. Biomaterials. 26:6375–6385. doi:10.1016/j.biomaterials.2005.03.043.

Klinman, D.M., A.K. Yi, S.L. Beaucage, J. Conover, and A.M. Krieg. 1996. CpG motifs present in bacteria DNA rapidly induce lymphocytes to secrete interleukin 6, interleukin 12, and interferon gamma. Proc. Natl. Acad. Sci. U. S. A. 93:2879–2883.

Leleux, J., and K. Roy. 2013. Micro and Nanoparticle-Based Delivery Systems for Vaccine Immunotherapy: An Immunological and Materials Perspective. Adv. Healthc. Mater. 2:72–94. doi:10.1002/adhm.201200268.

Liu, H., K.D. Moynihan, Y. Zheng, G.L. Szeto, A.V. Li, B. Huang, D.S. Van Egeren, C. Park, and D.J. Irvine. 2014. Structure-based programming of lymph-node targeting in molecular vaccines. Nature. 507:519–522. doi:10.1038/nature12978.

Lund, J., A. Sato, S. Akira, R. Medzhitov, and A. Iwasaki. 2003. Toll-like Receptor 9-mediated Recognition of Herpes Simplex Virus-2 by Plasmacytoid Dendritic Cells. J. Exp. Med. 198:513–520. doi:10.1084/jem.20030162.

Lynn, G.M., R. Laga, P.A. Darrah, A.S. Ishizuka, A.J. Balaci, A.E. Dulcey, M. Pechar, R. Pola, M.Y. Gerner, A. Yamamoto, C.R. Buechler, K.M. Quinn, M.G. Smelkinson, O. Vanek, R. Cawood, T. Hills, O. Vasalatiy, K. Kastenmüller, J.R. Francica, L. Stutts, J.K. Tom, K.A. Ryu, A.P. Esser-Kahn, T. Etrych, K.D. Fisher, L.W. Seymour, and R.A. Seder. 2015. In vivo characterization of the physicochemical properties of polymer-linked TLR agonists that enhance vaccine immunogenicity. Nat. Biotechnol. 33:1201–1210. doi:10.1038/nbt.3371.

Moon, J.J., H. Suh, A. Bershteyn, M.T. Stephan, H. Liu, B. Huang, M. Sohail, S. Luo, S. Ho Um, H. Khant, J.T. Goodwin, J. Ramos, W. Chiu, and D.J. Irvine. 2011. Interbilayer-crosslinked multilamellar vesicles as synthetic vaccines for potent humoral and cellular immune responses. Nat. Mater. 10:243–251. doi:10.1038/nmat2960.

Mothe, B.R., H. Horton, D.K. Carter, T.M. Allen, M.E. Liebl, P. Skinner, T.U. Vogel, S. Fuenger, K. Vielhuber, W. Rehrauer, N. Wilson, G. Franchini, J.D. Altman, A. Haase, L.J. Picker, D.B. Allison, and D.I. Watkins. 2002. Dominance of CD8 Responses Specific for Epitopes Bound by a Single Major Histocompatibility Complex Class I Molecule during the Acute Phase of Viral Infection. J. Virol. 76:875–884. doi:10.1128/JVI.76.2.875-884.2002.

Nefedova, Y., P. Cheng, D. Gilkes, M. Blaskovich, A.A. Beg, S.M. Sebti, and D.I. Gabrilovich. 2005. Activation of dendritic cells via inhibition of Jak2/STAT3 signaling. J. Immunol. Baltim. Md 1950. 175:4338–4346.

Ohto, U., T. Shibata, H. Tanji, H. Ishida, E. Krayukhina, S. Uchiyama, K. Miyake, and T. Shimizu. 2015. Structural basis of CpG and inhibitory DNA recognition by Toll-like receptor 9. Nature. 520:702–705. doi:10.1038/nature14138.

Pai Kasturi, S., H. Qin, K.S. Thomson, S. El-Bereir, S.-C. Cha, S. Neelapu, L.W. Kwak, and K. Roy. 2006. Prophylactic anti-tumor effects in a B cell lymphoma model with DNA vaccines delivered on polyethylenimine (PEI) functionalized PLGA microparticles. J. Control. Release Off. J. Control. Release Soc. 113:261–270. doi:10.1016/j.jconrel.2006.04.006.

Piccioli, D., C. Sammicheli, S. Tavarini, S. Nuti, E. Frigimelica, A.G.O. Manetti, A. Nuccitelli, S. Aprea, S. Valentini, E. Borgogni, A. Wack, and N.M. Valiante. 2009. Human plasmacytoid dendritic cells are unresponsive to bacterial stimulation and require a novel type of cooperation with myeloid dendritic cells for maturation. Blood. 113:4232–4239. doi:10.1182/blood-2008-10-186890.

Pradhan, P., H. Qin, J.A. Leleux, D. Gwak, I. Sakamaki, L.W. Kwak, and K. Roy. 2014. The effect of combined IL10 siRNA and CpG ODN as pathogen-mimicking microparticles on Th1/Th2 cytokine balance in dendritic cells and protective immunity against B cell lymphoma. Biomaterials. 35:5491–5504. doi:10.1016/j.biomaterials.2014.03.039.

Pulendran, B., J.L. Smith, G. Caspary, K. Brasel, D. Pettit, E. Maraskovsky, and C.R. Maliszewski. 1999. Distinct dendritic cell subsets differentially regulate the class of immune response in vivo. Proc. Natl. Acad. Sci. 96:1036–1041. doi:10.1073/pnas.96.3.1036.

Reddy, S.T., A. Rehor, H.G. Schmoekel, J.A. Hubbell, and M.A. Swartz. 2006. In vivo targeting of dendritic cells in lymph nodes with poly(propylene sulfide) nanoparticles. J. Controlled Release. 112:26–34. doi:10.1016/j.jconrel.2006.01.006.

Samarasinghe, R., P. Tailor, T. Tamura, T. Kaisho, S. Akira, and K. Ozato. 2006. Induction of an anti-inflammatory cytokine, IL-10, in dendritic cells after toll-like receptor signaling. J. Interferon Cytokine Res. Off. J. Int. Soc. Interferon Cytokine Res. 26:893–900. doi:10.1089/jir.2006.26.893.

Saraiva, M., and A. O’Garra. 2010. The regulation of IL-10 production by immune cells. Nat. Rev. Immunol. 10:170–181. doi:10.1038/nri2711.

Singh, A., H. Nie, B. Ghosn, H. Qin, L.W. Kwak, and K. Roy. 2008. Efficient modulation of T-cell response by dual-mode, single-carrier delivery of cytokine-targeted siRNA and DNA vaccine to antigen-presenting cells. Mol. Ther. J. Am. Soc. Gene Ther. 16:2011–2021. doi:10.1038/mt.2008.206.

Singh, A., H. Qin, I. Fernandez, J. Wei, J. Lin, L.W. Kwak, and K. Roy. 2011. An injectable synthetic immune-priming center mediates efficient T-cell class switching and T-helper 1 response against B cell lymphoma. J. Control. Release Off. J. Control. Release Soc. 155:184–192. doi:10.1016/j.jconrel.2011.06.008.

Singh, A., S. Suri, and K. Roy. 2009. In-situ crosslinking hydrogels for combinatorial delivery of chemokines and siRNA-DNA carrying microparticles to dendritic cells. Biomaterials. 30:5187–5200. doi:10.1016/j.biomaterials.2009.06.001.

Stano, A., C. Nembrini, M.A. Swartz, J.A. Hubbell, and E. Simeoni. 2012. Nanoparticle size influences the magnitude and quality of mucosal immune responses after intranasal immunization. Vaccine. 30:7541–7546. doi:10.1016/j.vaccine.2012.10.050.

Swartz, M.A., J.A. Hubbell, and S.T. Reddy. 2008. Lymphatic drainage function and its immunological implications: from dendritic cell homing to vaccine design. Semin. Immunol. 20:147–156. doi:10.1016/j.smim.2007.11.007.

Vogt, P.K., and J.R. Hart. 2011. PI3K and STAT3: A New Alliance. Cancer Discov. 1:481–486. doi:10.1158/2159-8290.CD-11-0218.

